# Locality sensitive hashing for the edit distance

**DOI:** 10.1101/534446

**Authors:** Guillaume Marçais, Dan DeBlasio, Prashant Pandey, Carl Kingsford

## Abstract

**Motivation:** Sequence alignment is a central operation in bioinformatics pipeline and, despite many improvements, remains a computationally challenging problem. Locality Sensitive Hashing (LSH) is one method used to estimate the likelihood of two sequences to have a proper alignment. Using an LSH, it is possible to separate, with high probability and relatively low computation, the pairs of sequences that do not have an alignment from those that may have an alignment. Therefore, an LSH reduces in the overall computational requirement while not introducing many false negatives (i.e., omitting to report a valid alignment). However, current LSH methods treat sequences as a bag of *k*-mers and do not take into account the relative ordering of *k*-mers in sequences. And due to the lack of a practical LSH method for edit distance, in practice, LSH methods for Jaccard similarity or Hamming distance are used as a proxy.

**Results:** We present an LSH method, called Order Min Hash (OMH), for the edit distance. This method is a refinement of the minHash LSH used to approximate the Jaccard similarity, in that OMH is not only sensitive to the *k*-mer contents of the sequences but also to the relative order of the *k*-mers in the sequences. We present theoretical guarantees of the OMH as a gapped LSH.

## 1 Introduction

Measuring sequence similarity is the core of many algorithms in computational biology. For example, in the Overlap-Layout-Consensus paradigm to assemble genomes (e.g., Myers *et al.*, 2000; Jaffe *et al.*, 2003), the first overlap step consist of aligning the reads against one another to determine which pairs have a significant alignment (an overlap). In meta-genomics, sequencing reads, or longer sequences created from these reads, are aligned against known genomes, or against one another to cluster the sequences, to determine the constituent species of the sample. Sequence similartiy is also at the heart of the many general sequence aligners, either genome to genome (e.g., MUMmer Marçais *et al.*, 2018) or reads to genome [e.g., Bowtie2 (Langmead and Salzberg, 2012), BWA (Li and Durbin, 2010)], that are used in countless pipelines in bioinformatics

Despite many algorithmic and engineering improvements [e.g., implementation on SIMD (Zhao *et al.*, 2013) and GPU (Liu *et al.*, 2012)], computing the sequence alignment or edit distance between two sequences takes approximately quadratic time in the length of the input sequences, which remains computationally expensive in practice. Given that the edit distance is likely not computable in strong subquadratic time (Backurs and Indyk, 2015), most aligners rely on heuristics to more quickly detect sequences with a high probability of having an alignment.

Recent aligners, such as Mash (Ondov *et al.*, 2016), Mashmap (Jain *et al.*, 2017), or overlappers such as MHap (Berlin *et al.*, 2015), use a method called “Locality Sensitive Hashing” (LSH) to reduce the amount of work necessary (Indyk and Motwani, 1998). The procedure is a dimensionality reduction method and works in two steps. First, the sequences (or part of the sequences) are summarized into *sketches* that are much smaller than the original sequences while preserving important information to estimate how similar two sequences are. Second, by directly comparing those sketches (with no need to refer to the original sequences) or by using these sketches as keys into hash tables, the software finds pairs of sequences that are likely to be similar. A more thorough, and computationally expensive, alignment procedure may then be used on the candidate pairs to refine the actual alignments.

In an LSH method, the distance between sketches is used as a first approximation for the distance between the sequences. That is, with high probability, two sequences which are very similar must have sketches which are similar, and conversely dissimilar sequences have dissimilar sketches. More precise definition of these concepts are given in Section 2.

Instead of using an LSH for the edit distance or an alignment score, in practice sequence alignment programs use the minHash LSH (Broder, 1997) for the Jaccard similarity or an LSH for the Hamming distance as a proxy for the edit distance. While these two techniques have proven themselves useful in practice, they suffer from one major flaw: neither the Jaccard similarity nor the Hamming similarity directly correspond to the edit distance (see Section 2.2 for examples). In fact, it is possible to find sequences that are indistinguishable according to the Jaccard similarity, but have large edit distance. Similarly, with the Hamming distance, there exists sequences with very low edit distance that are completely dissimilar according to the Hamming similarity.

Depending on the problem and the software implementation, the cases above can lead to false negatives (an alignment is missed) and a decrease in precision, or false positives (a nonexistent potential alignment reported) and extra computational work. An LSH method for edit distance instead of the proxy Jaccard or Hamming similarities would reduce both of these issues.

Although multiple definitions are possible for sequence similarity (or distance), in this study, we focus on the edit distance (aka Levenshtein distance, Levenshtein, 1966), which is the number of operations (misma tches, insertion, deletion) needed to transform a string into another one.

Two methods that are LSH for the edit distance have been described previously. Bar-Yossef *et al.* (2004) propose a sketch that can distinguish, with some probability, between sequences with edit distance ≤ *t* from sequences with edit distance ≥ (*tn*)^2/3^, where *n* is the length of the sequencesm, for any 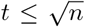. They use an indirect method to obtain an LSH for the edit distance: first they embed the edit distance space into a Hamming space with low distortion, and second, apply an LSH on the Hamming space. That is, the input sequence is first transformed into a bit vector of high dimension, then sketching for the Hamming distance is applied to obtain an LSH for the edit distance.

Similarly, Ostrovsky and Rabani (2007) propose a two step method, where the edit distance space is first embedded into an *ℓ*_1_ space with low distortion, then a sketching algorithm for the *ℓ*_1_ (Kushilevitz *et al.*, 2000) is used to obtain an LSH for the edit distance. This method can distinguish between sequences with edit distance ≤ *t* and edit distance 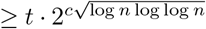 for some constant *c*.

We propose a simpler and direct method that is an LSH for the edit distance. Our method is an extension to the minHash method. We call our method OMH for Order Min Hash, and it can be seen as a correction of the minHash method. The probability of hash collision in the OMH method is the product of two probabilities. The first is the probability to select a *k*-mer from the set of common *k*-mers between the two sequences. This probability is similar to minHash that estimates the Jaccard similarity between the *k*-mer contents of two sequences. However, there is one key difference: the minHash method estimates the Jaccard similarity which treats sequences as sets of *k*-mers, and the number of occurrences of each *k*-mer in the sequences is ignored. Whereas OMH estimates the *weighted Jaccard*, where the number of occurrences of a *k*-mer in a sequence is significant, i.e., the weighted Jaccard works with multi-sets. The second probability is the likelihood that the common *k*-mers appear in the same relative order in the two sequences. Therefore, OMH is not only sensitive to the *k*-mer content of the sequences but also to the order of the *k*-mers in the sequences.

The sketch proposed for OMH is only slightly bigger than the sketch for minHash while maintaining significantly more information about the similarity of two sequences. In addition to providing an estimate for the edit distance between two sequences, it also provides an estimate of the *k*-mer content similarity (the weighted Jaccard) and how similar the relative order is between the common *k*-mers of the two sequences.

Section 2 summarizes the notation used though out and main results. Detailed proofs of the results are given in Section 3. Section 4 discusses some practical consideration on the implementation of the sketches.

## 2 Main results

### 2.1 Concepts and definitions

#### Distance and similarity

A *distance* is a function 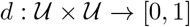 that indicates the distance between two elements in the universe 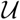. *d* satisfies the triangle inequality and *d*(*x*, *y*) = 0 means that *x* = *y*. A *similarity* is a function *s*(·, ·) such that 1−*s* is the distance. Hence, a distance defines a similarity and vice-versa. We will therefore use equivalently the terms “edit distance” and “edit similarity”.

Given two strings *S*_1_, *S*_2_ ∈ Σ^*n*^ of length *n* (where Σ is the alphabet of size *σ* = |Σ|) Hamming distance H_d_(*S*_1_, *S*_2_) is the number of indices at which *S*_1_ and *S*_2_ differ divided by *n*: H_d_(*S*_1_, *S*_2_) = |{*i* ∈ [*n*] | *S*_1_ [*i*] ≠ *S*_2_ [*i*]|/*n* ([*n*] denotes the set (0,…,*n* − 1}). The (normalized) edit distance E_d_(*S*_1_, *S*_2_) is the minimum number of indels (sort for insertion or deletion), and mismatches necessary to transform *S*_1_ into *S*_2_ divided by *n*. Given two sets *A* and *B*, the Jaccard similarity is J(*A*, *B*) = |*A* ∩ *B*|/|*A* ∪ *B*|.

#### Gapped LSH

Let 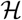 be a set of hash functions defined on a set 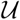 (the universe). A probability distribution on the set 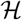 is called (*s*_1_, *s*_2_, *p*_1_, *p*_2_)-*sensitive* for the similarity *s* when

- 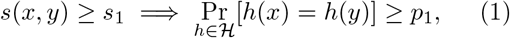
- 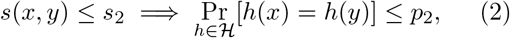

where *s*_1_ ≥ *s*_2_ and *p*_1_ ≥ *p*_2_. A similarity admits a *gapped LSH* scheme if there exists a distribution on a set of hash functions that is *s*_1_, *s*_2_, *p*_1_, *p*_2_)-sensitive. In the definition above, the probability is taken over the choice of the hash function in 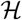 and the implications hold for any choice of *x* and 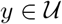 In a gapped LSH, the probability of a hash collision is increased (≥ *p*_1_) between similar elements, and less likely (≤ *p*_2_) for dissimilar elements.

In the following, the probabilities are always taken over the choice of the hashing function, even though we may omit the 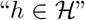 subscript.

#### LSH

An LSH for a similarity is a family of hash functions that is (*r*, *r*, *r*, *r*)-sensitive for any *r* ∈ (0, 1). Equivalently, the family of hash functions satisfies Pr[*h*(*x*) = *h*(*y*) = *s*(*x*, *y*). In practice a gapped LSH is typically used to put elements into a hash table where there is high likelyhood of a collision, while a full LSH can be used as a direct estimator of the underlying measurement.

#### MinHash sketch

Let the universe 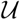 be a family of sets on the ground set *X* (i.e., 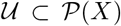). The minHash LSH for the Jaccard similarity is defined as the uniform distribution on the set 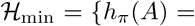 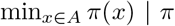 is a permutation of *X*}. That is, the hash function selects the smallest element of the set *A* according to some ordering *π* of the elements of the ground set *X*. This family of hash functions is (*s*, *s*, *s*, *s*)-sensitive for any value of *s* ∈ [0, 1], or equivalently Pr[*h*(*A*) = *h*(*B*)] = J(*A*, *B*).

#### LSH for Hamming similarity

The Hamming similarity between two sequences is the proportion of positions which are equal: H_s_(*S*_1_, *S*_2_) = |{*i* ∈ [*n*] | *S*_1_[*i*] = *S*_1_[*i*]}|/*n*. For the Hamming similarity, the uniform distribution on 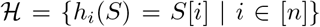 satisfies Pr[*h*(*S*_1_) = *h*(*S*_2_)] = H_s_(*S*_1_, *S*_2_).

#### String *k*-mer set

For a sequence *S*, the set of its constituent *k*-mers is 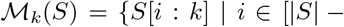 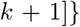 where *S*[*i*: *k*] is the substring of length *k* starting at index *i*. By extension, the Jaccard between two sequences is the Jaccard between their *k*-mer sets: 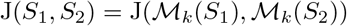

#### Weighted Jaccard

The weighted Jaccard similarity on multisets (or weighted sets) is defined similarly to the Jaccard similarity on sets, where the intersection and union take the multiplicity of the elements into account. More precisely, a multiset *A* is defined by an index function 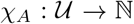, where χ_*A*_(*x*) gives the multiplicity of *x* in *A* (zero if not present in *A*). The index function of the intersection of two multisets is the minimum of the index functions, and for the union it is the maximum. Then, the weighted Jaccard is defined by

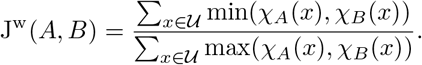

This is a direct extension to the set definitions, where the index function takes values in {0, 1}.

### 2.2 Jaccard and Hamming differ from edit distance

Similarly to the definition of the LSH, we say that a similarity f1 is a (*s*_1_, *s*_2_, *t*_1_, *t*_2_)-proxy for the similarity *f*_2_ if

- 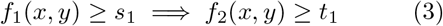
- 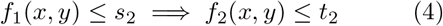

That is, high similarity for *f*_1_ implies high similarity for *f*_2_, and the converse. Because of the symmetry in the definitions between sensitivity and proxy, if *f*_1_ is not a proxy for *f*_2_, then an LSH for *f*_1_ is not an LSH for *f*_2_.

We show here that neither the Hamming similarity nor the Jaccard similarity are good proxies for the edit distance. More precisely, only one of the implications above is satisfied.

#### Jaccard similarity differs from edit similarity

A low Jaccard similarity does imply a low edit similarity (eq (4)). On the other hand, consider the sequence *S*_1_ = 0…01…1 that has *n* − *k* 0s followed by *k* 1s, and *S*_2_ with *k* 0s followed by *n* − *k* 1s (*k* fixed, *n* arbitrarily large). The *k*-mer sets of *S*_1_ and *S*_2_ are identical, hence J(*S*_1_, *S*_2_) = 1, while the edit similarity is ≤ 2*k*/*n*. These sequence are indistinguishable according to the Jaccard similarity while having arbitrarily small edit similarity (eq (3) not satisfied).

#### Weighted Jaccard similarity differs from edit similarity

Consider two de Bruijn sequences: sequences of length *σ*^*k*^ containing every *k*-mer exactly once (van Aardenne-Ehrenfest and de Bruijn, 1951). There is a very, very large number of such sequences 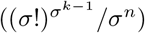, and although any two such sequences have exactly the same *k*-mer content, they might otherwise have a very low edit similarity. Both the Jaccard and weighted Jaccard similarity fail to distinguish between de Bruijn sequences, regardless of their mutual edit distance.

More generally, both Jaccard and weighted Jaccard similarity treats sequences as bags of *k*-mers. The information on relative order of these *k*-mers within the sequence is ignored, although it is of great importance for the edit similarity. By contrast, an OMH sketch does retain some information on the order of the *k*-mers in the original sequence.

#### Hamming similarity differs from edit similarity

A high Hamming similarity does imply a high edit similarity (eq (3)). The opposite is not true however. Consider the sequences of length *n*, *S*_1_ = 0101…01 and *S*_2_ = 1010…10. These sequences have a Hamming similarity of 0 and an edit similarity of ≥1−2/*n* (two indels). That is, these sequences are as dissimilar as possible according to the Hamming distance, but an arbitrarily high edit similarity (eq (4) not satisfied).

The Hamming similarity is very sensitive to the absolute position in the string. A single shift between two sequence has a large impact on the Hamming similarity but only a unit cost for the edit similarity. An OMH sketch on the other hand only contains relative order between *k*-mers and is indifferent to changes in absolute position.

### 2.3 LSH for the edit similarity

An LSH for the edit similarity must be sensitive to the *k*-mer content of the strings and the relative order these *k*-mers, but relatively insensitive to the absolute position of the *k*-mers in the string. This motivates the definition below. Similarly to the minHash, *k*-mers are selected at random by using a permutation on the *k*-mers. Additionally, to preserve information about relative order, **ℓ* k*-mers are selected at once and recorded in the order they appear in the sequence (rather than the order defined by the permutation).

Additionally, the method must handle repeated *k*-mers. Two copies of the same *k*-mer occur at different positions in the sequence, and it is important for the relative ordering between *k*-mers to distinguish between these two copies. We make *k*-mers unique by appending to them their “occurrence number”.

More precisely, for a string *S* of length |*S*| = *n*, consider the set 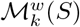 of the pairs of the *k*-mers and their occurrence number. If there are *x* copies of *m* in sequence *S*, then the *x* pairs (*m*, 0),…,(*m*, *x* − 1) are in the set 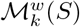, and the occurrence number denotes the number of other copies of *m* that are in the sequence *S* to the left of this particular copy. That is, if *m* is the *k*-mer at position *i* in *S* (i.e., *m* = *S*[*i*: *k*]), then its occurrence number is |{*j* ∈ [*i*] | *S*[*j*: *k*] = *m*}|. This set is the “multi-set” of the *k*-mer content of string *S*, or the “weighted set” of *k*-mers where the number of occurrences is the weight of the *k*-mer (hence the *w* superscript). We call a pair (*m*, *i*) of a *k*-mer and an occurrence number a “uniquified” *k*-mer.

A permutation π of ∑^*k*^ × [*n*] defines two functions 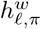 and *h*_**ℓ**,*π*_. 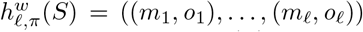 is a vector of length **ℓ** of elements of 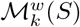 such that:

- the pairs (*m*_*i*_, *o*_*i*_) are the **ℓ** smallest elements of 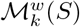 according to *π*,
- the pairs are listed in the vector in the order in which the *k*-mer appear in the sequence *S*. That is, if *i* < *j*, *m*_*i*_ = *S*[*x*: *k*] and *m*_*j*_ = *S*[*y*: *k*], then *x* < *y*.

The vector *h*_**ℓ**,*π*_(*S*) = (*m*_1_,…,*m*_**ℓ**_) contains only the *k*-mers from 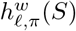, in the same order. The Order Min Hash method (OMH) is defined as the uniform distribution on the set of hash function 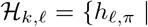 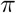 a permutation of ∑^*k*^ × [*n*]}.

For extreme cases, where **ℓ** = *n* − *k* + 1, the vector contains overlapping *k*-mers that cover the entire sequence *S*. In that case, equality of the hash values implies strict equality of the sequences.

At the other extreme, where **ℓ** = 1, the vectors contain only one *k*-mer and no relative order information is preserved. In that case, only the *k*-mer content similarity between *S*_1_ and *S*_2_ matters. Define the *weighted Jaccard* similarity as 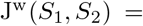 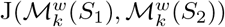.

#### Theorem 1.

When ℓ = 1, OMH is an LSH for the weighted Jaccard similarity:

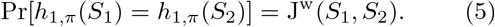

#### Proof.

This proof is similar to that of minHash and the Jaccard similarity (Broder, 1997). Because every uniquified *k*-mer in 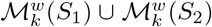 has the same probability of being selected, the probability of having a hash collision is the same as selected a *k*-mer from the intersection where the probability of picking a *k*-mer is weighted by its maximum occurrence number.

As we shall see in Section 4.4, the weighted Jaccard similarity contains approximately the same information as the Jaccard similarity with respect to the edit similarity.

For the general case 1 < **ℓ** < *n* − *k* + 1, we shall prove the following theorem in Section 3 that OMH is a gapped LSH for the edit distance.

#### Theorem 2.

There exists functions 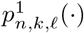 and 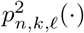 such that OMH is 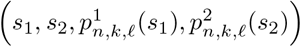-sensitive for any ℓ ∈ [2, n − k].

The actual functions *p*^1^ and *p*^2^ are explicitly define in Section 3, but they may not be easily expressed with elementary functions in general.

## 3 Proofs of main results

We shall now prove Theorem 2 that OMH is sensitive for the edit similarity by exhibiting the relations between parameters (*s*_1_, *s*_2_, *p*_1_, *p*_2_) that satisfy eq (1) and (2). We will break the proof in two Lemmas that provide the relations between *s*_1_, *p*_1_ and between *s*_2_, *p*_2_. In the following, *S*_1_ and *S*_2_ are two sequences of length *n*. The number of *k*-mers in each of these sequences is *n*_*k*_ = *n* − *k* + 1.

### Lemma 1.

E_s_(S_1_,S_2_) ≥ s_1_ ⟹ Pr[h_ℓ,π_(S_1_) = h_ℓ,π_(S_2_)]≥ p_1_ when

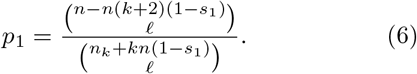

### Proof.

The situation is similar to the minHash method. Suppose that E_s_(*S*_1_, *S*_2_) ≥ *s*_1_, then the edit distance E_d_(*S*_1_, *S*_2_) = 1 − E_s_(*S*_1_, *S*_2_) ≤ 1 − *s*_1_ and the number of mismatch and indels is ≤ *n*(1 − *s*_1_). At least *n*_*k*_ − *kn*(1 − *s*_1_) *k*-mers must be part of an optimal alignment as an error (mismatch or indel) affects at most *k* consecutive *k*-mers.

Similarly, the size of the set 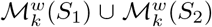 is maximized when all the *k*-mers that are not part of the alignment are different. Then 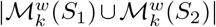 is at most *n*_*k*_ + *kn*(1 − *s*_1_).

We estimate the probability to have a hash collision from the number of uniquified *k*-mers in the alignment. As seen in Figure 1, it is possible for a *k*-mer *m* with different occurrence numbers to be part of the optimal alignment. Any permutation that has (*m*, 0) in the lowest **ℓ** uniquified *k*-mers does not lead to a hash collision. Because for any such uniquified *k*-mer (*m*, 0) in the alignment, there is also an instance in the unaligned part of *S*_1_ and *S*_2_, the number *x* of such *k*-mers satisfy *x* + *k* − 1 ≤ 2*n*(1 − *s*_1_). That is, the total sequence covered by these *k*-mers cannot exceed the total unaligned sequence between *S*_1_ and *S*_2_. Consequently, the number of *k*-mers in the alignment to choose from is at least *n* − *n*(*k* + 2)(1 − *s*_1_).

**Figure 1:**
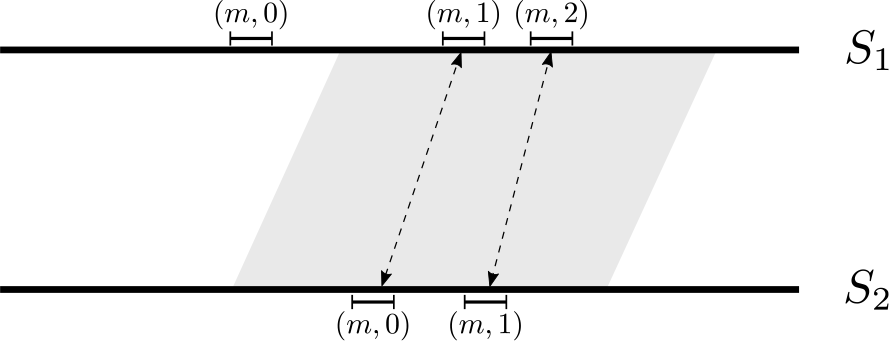
The gray area represents an optimal alignment between sequence *S*_1_ and *S*_2_. A particular *k*-mer *m* is shown with its occurrence numbers. The occurrence number of the matched *k*-mer pairs in the alignment may not agree (as in this example). For every such *m* with a mismatch occurrence number in
the alignment, there must exists an instance of *m* not contained in the alignment ((*m*, 0) in *S*_1_ here).

Every element of 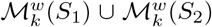 has an equal probability to be in the lowest **ℓ** elements according to a permutation *π*, therefore the probability of having a hash collision is:

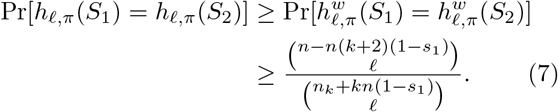

This defines the relationship between *p*_1_ and *s*_1_ as in eq (6).

For the proof of eq (2), we will consider the contrapositive

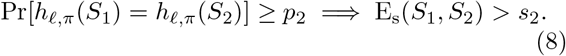

That is for any two sequences with high probability of having a hash collision, the edit similarity of the sequences must be high.

To have a high probability of collision between two sketches, the sequences must have a large number of common *k*-mers and these common *k*-mers should be mostly in the same relative order. The first condition corresponds to the sequences having a large weighted Jaccard similarity.

The second condition is related to common subsequences. A “common subsequence” between (*a*_*i*_) and (*b*_*i*_) is a sequence of elements that are in both (*a*_*i*_) and (*b*_*i*_) and appear in the same order (formally an increasing function such that *φ* such that *a*_*φ*(*i*)_ = *b*_*φ*(*i*)_). If the sequences *S*_1_ and *S*_2_ have a long common subsequence of *k*-mers, then the probability to pick **ℓ* k*-mers in the same order between the common *k*-mers of *S*_1_ and *S*_2_ will be high. In turn the presence of a long common subsequence of *k*-mers must have a high similarity.

Hence, the proof will rely on finding pairs of sequences with as short as possible longest common subsequences while having high probability to pick *k*-mers in the same order. This problem is equivalent to finding, for a given *L*, a single sequence with as many as possible increasing subsequences of length *L* (see Lemma 3).

### Lemma 2.

E_s_(S_1_, S_2_) ≤ s_2_ ⟹ Pr[h_ℓ,π_(S_1_) = h_ℓ,π_(S_2_)] ≥ p_2_ when

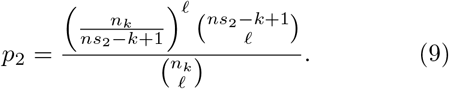

### Proof.

We use the notation 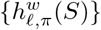 for the set of elements in the vector 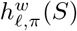 (in other words, because all the elements are unique by construction, the elements without order).

As mentioned above, we consider the contrapositive state in eq (8). We have that

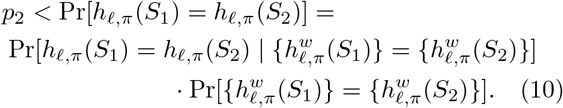

Under the conditional event (*C*) that 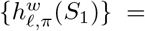 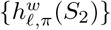, we have 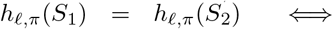 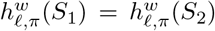. The reverse implication (⇐) always true as *h* is obtained from *h*^*w*^ by using only the *k*-mer in each element. The forward implication (⇒) holds thanks to (*C*). Given that the *k*-mers are listed in order in which they appear in the respective sequences, they are also listed in order of their occurrence number, and because the content in the weighted vectors *h*^*w*^ is the same, the equality of the unweighted vectors *h* implies equality of the weighted vectors.

Let 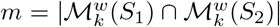 be the size of the intersection of the weighted *k*-mer sets. The event (*C*) occurs when the *ℓ* smallest according to the permutation *π* belong to the intersection 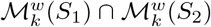.

Therefore

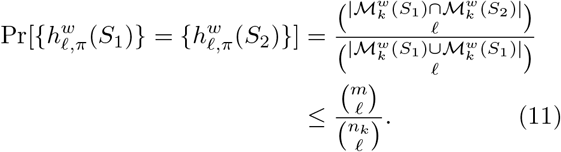

Consider now the sequences 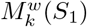 and 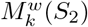 of the elements of 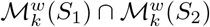 listed in the order in which their occur in *S*_1_ and *S*_2_ respectively. Both of these sequences have length *m*. Then, the event that *h*_*ℓ*,*π*_ = *h*_*ℓ*,*π*_(*S*_2_) under the condition (*C*) is equivalent to having the hash function 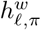 picking a common subsequence (CS) of length *ℓ* between 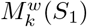 and 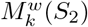. Because the elements of these sequences are never repeated (it is a list of uniquified *k*-mers), the problem of finding common subsequences between 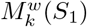 and 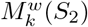 is identical to finding increasing subsequences (IS) in a sequence of integers of length *m* (Hunt and Szymanski, 1977; Fredman, 1975).

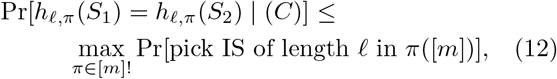

where [*m*]! is the set of all permutations of [*m*]. Together, equations (10), (11), (12), and lemma 3 below imply that the following holds for any choice of sequences *S*_1_*, S*_2_:

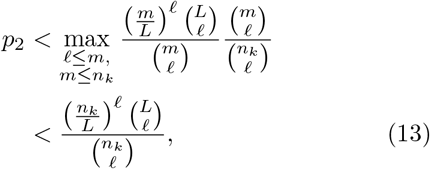

where *L* is the length of the longest common subsequence between 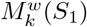 and 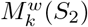. The function on the right hand side of eq (13) is an increasing function of *L*, equal to 0 when *L* < *ℓ*, and equal to 1 when *L* = *n* − *k* + 1. Given that *s*_2_ ≥ *E*_*s*_ (*S*_1_, *S*_2_) ≥ (*L* + *k* − 1)/*n*, replacing *L* by *ns*_2_ − *k* + 1 in eq (13) gives the desired relation between *s*_2_ and *p*_2_ of eq (9).

Finally, we prove the relationship between the length of the longest increasing subsequence and the largest number of sequences of maximal length.

### Lemma 3.

For i, n, ℓ ∈ ℕ, n ≥ i ≥ ℓ, for any sequence of length n with a longest increasing subsequence (LIS) of at most i, the largest number of increasing subsequences of length ℓ is

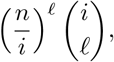

and this bound is tight.

### Proof.

The proof relies on the properties of patience sorting (Aldous and Diaconis, 1999). Patience sorting for a shuffled deck of cards works as follows:

- The algorithm creates stacks of cards where in each stack the cards are in decreasing order from the bottom to the top of the stack. The stacks are organized in a line, left to right.
- At each round, the next card of the deck is examined and added to the top of the left most stack it can go on, i.e., the left most stack with a top card whose value is higher that the new card.
- If no existing stack is suitable, a new stack is created to the right with the new card.

After all the cards are drawn and organized in stacks (see Figure 2), the following properties hold: (1) no two cards from an increasing subsequence in the original deck are in the same stack, and (2) the number of stacks is equal to the LIS (see Aldous and Diaconis (1999, Lemma 1)).

**Figure 2:**
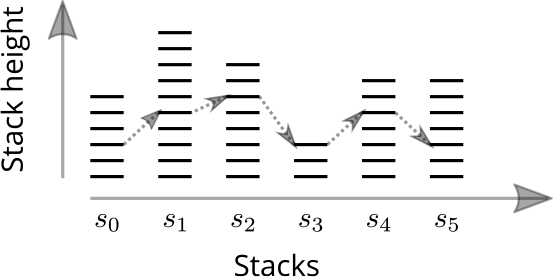
Figure 2: An example of stacks created when sorting a deck of cards. The LIS is 6, with the arrows showing a possible increasing subsequence of maximal length. To maximize the number of possible subsequences of maximum length, the height of the stacks have to be equal.

Fix *i*′ ∈ [*ℓ*, *i*] and a sequence *S* of length *n* with LIS of *i*′. At the end of patience sorting of *S*, let **s** = (*s*_0_,…, *s*_*i*′ − 1_) be the vector of the height of each of the stacks. Then, an upper bound on the number of increasing subsequence of length *ℓ* in *S* is

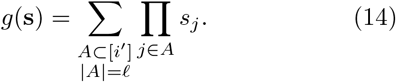

This is an upper bound as every choice of *ℓ* elements from different stacks does not necessarily define a valid increasing subsequence of *S*. We show that *g* reaches its maximum when *s*_0_ =…= *s*_*i*′ − 1_ = *n*/*i*′.

Because the set *C* = {**s** = (*s*_0_,…, *s*_*i*′_) | ∑_*j*_ *s*_*j*_ = *n* is compact, *g* reaches a maximum on *C*. Suppose that in **s**, not all the *s*_*j*_ are equal; WLOG assume that *s*_*i*′ − 2_ and *s*_*i′ − 1*_ are distinct. Set *α* = (*s*_*i*′ − 2_ + *s*_*i*′ − 1_)/2 and consider the point **s**′ = (*s*_0_,…, *s*_*i*′ − 3_, *α, α*). Let us also use the notation

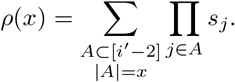

Then, we split the sum in *g*(**s**′) into the terms containing neither *s*_*i*′ − 2_ nor *s*_*i*′ 1_ (= *ρ*(*ℓ*)), the terms that contain one of *s*_*i*′ − 2_ or *s*_*i*′ − 1_ (= 2*αρ*(*ℓ* − 1)) and the terms that contain both (= *α*^2^*ρ*(*ℓ* − 1)), and we use the inequality *α*^2^ > *s*_*i*′ − 2_*s*_*i*′ − 1_ (arithmetic mean is larger than geometric mean):

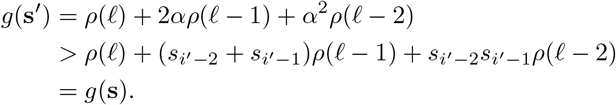

Hence, *g*(**s**), where **s** contains two distinct values, is not maximum, and *g* must reach its maximum when all the *s*_*j*_ are equal. Furthermore, in that case *s*_*j*_ = *n*/*i*′.

Therefore,

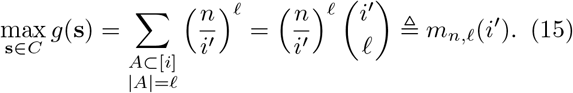

The function *m*_*n*,*ℓ*_(·) defined above is increasing and the maximum is reached for *i*′ = *i*.

Finally, consider the sequence *S*(*i, n*), *i* divides *n*, defined by blocks:

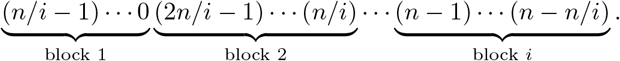

Each block is of length *n/i*, the numbers in each blocks are in decreasing order, and the start of the blocks are in increasing order: block *j* is the decreasing sequence (*jn*/*i*− 1)…((*j* − 1)*n*/*i*).

When the patience sorting algorithm is applied to the list *S*(*i, n*), the stacks are filled up one by one, from bottom to top and from left to right, and have the same height of *n/i*. Therefore, any choice of 1 element in each stack is a valid increasing subsequence of *S*(*i, n*) and the bound of equation (15) is attained.

Finally, we can conclude with the main theorem.

### Theorem 2.

*There exists functions* 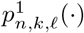 *and* 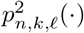 *such that OMH is* 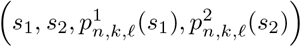-*sensitive for any ℓ* ∈ [2, *n* − *k*].

### Proof.

It is a direct consequence of Lemma 1 and Lemma 2.

## 4 Discussion

### 4.1 Parameters *s*_1_, *p*_1_, *s*_2_, *p*_2_

To have a proper LSH method, the conditions *p*_1_ ≥ *p*_2_ must hold. This condition means that the method is able to distinguish with some probability between dissimilar (≤ *s*_2_) and similar (≤ *s*_1_) sequences. Figure 3 shows the functions *p*^1^ (blue lines) and *p*^2^ (red lines) from Theorem 2 for varying values of *ℓ*.

**Figure 3.**
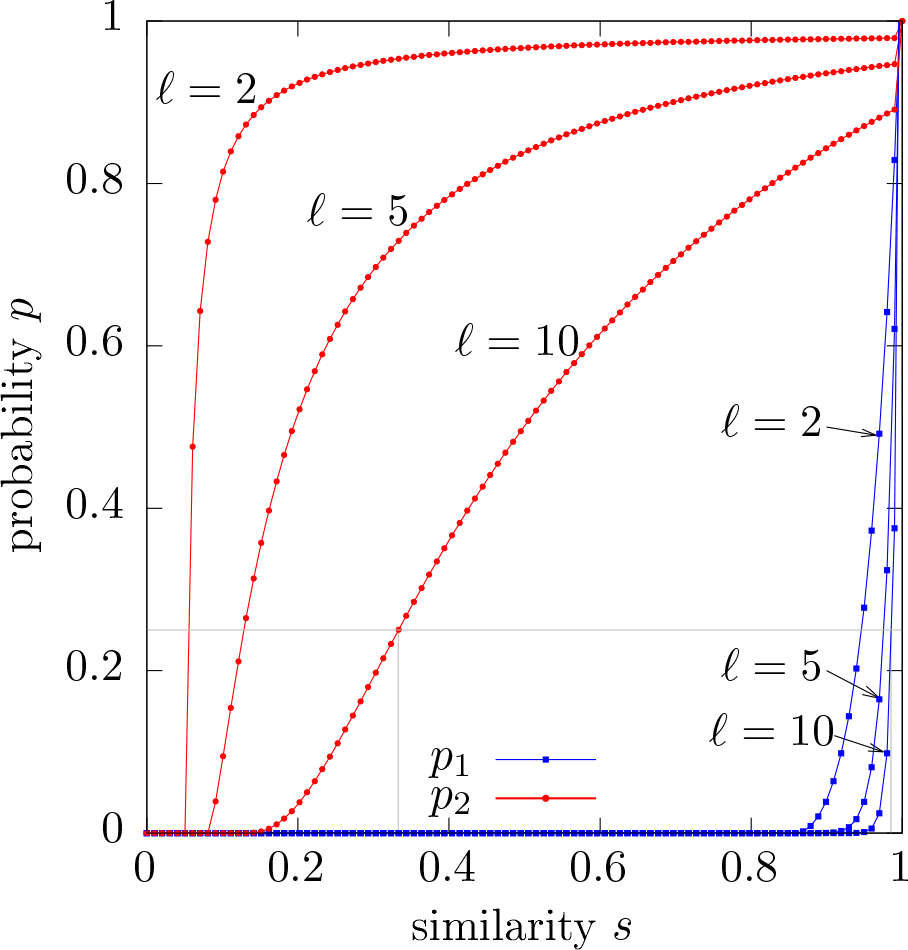
The relationships between the similarity thresholds *s*_1_ and *s*_2_, and the probabilities *p*_1_ and *p*_2_, for *n* = 100, *k* = 5. Given a probability, e.g., *p*_1_ = *p*_2_ = 0.25 shown by the horizontal gray line, the OMH method can distinguish between similarities below *s*_2_ (below left vertical gray line) and from similarities above *s*_1_ (above the right vertical gray line). The functions are defined at discrete points, when *ns* is integral, represented by the circles and squares.

At the limit, taking *p*_1_ = *p*_2_ = *p*, the method can distinguish between any *s*_1_ and *s*_2_ such that *p*^1^(*s*_1_) ≥ *p* and *p*^2^(*s*_2_) ≤ *p* (gray lines on Figure 3). For larger values of *ℓ*, the gap between distinguishable values is reduced, although at the cost of having high values for *s*_1_.

### 4.2 Choice of parameter *l*

The main difference between OMH and the minHash method is the choice of *ℓ k*-mers, where minHash correspond to the case of *ℓ* = 1 (ignoring the slight difference between Jaccard and weighted Jaccard). It might seem surprising at first that OMH is an LSH for edit distance for any values of *ℓ*, except for the extremes of *ℓ* = 1 and *ℓ* = *n* − *k* + 1.

The proof of Theorem 2 is consistent with this analysis. For both these extreme values of *ℓ*, eq (13), which relates the probability *p*_2_ to the similarity *s*_2_, becomes trivially true (*p*_2_ < 1). This means that even a certain hash collision (probability of 1) provides no guarantee on the relative order of common *k*-mers between the two sequences (i.e., the length of the longest subsequence *L* = 1: only 1 *k*-mer is guaranteed to align). On the other hand, any other value of *ℓ* leads to an actual bound in eq (13).

For example, when *ℓ* = 2, the minimum number of *k*-mers that must align in proper order as a function of the collision probability *p*_2_ is

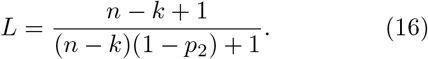

Even in the case where *n* is very large, then *L* ≈ 1/(1 − *p*_2_), the number of properly aligning *k*-mers becomes large when the probability of collision *p*_2_ is close to 1. This is in contrast to the minHash, where a probability of collision of 1 (i.e., a Jaccard similarity of 1) does not guarantee that more than 2 *k*-mers properly align (see example in Section 2.2).

In practice, the parameter *ℓ* should be relatively small, say 2 ≤ *ℓ* ≤ 5. Increasing the value of *ℓ* has two effects on the OMH method. First, it increases the minimum edit similarity that is detectable by the method, as there must be at least *ℓ* + *k* − 1 bases in the alignment of the two sequences for OMH to have a non-zero probability of hash collision. Second, a larger value of *ℓ* implies that the probability of hash collision is small, which requires storing a higher number of vectors in a sketch to obtain a low variance. There is a trade-off between how sensitive the scheme is to relative order (high value of *ℓ*) and the smaller size for the sketch (low value of *ℓ*).

### 4.3 Practical sketches for OMH

In our implementation, the OMH sketch for a sequence *S* contains more than just the list of vectors *h*_*ℓ*,*π*_(*S*).

In practice, we store

- the length of the sequence |*S*|,
- a list of *m* vectors *h*_*ℓ*,*π*_(*S*) and associated order vector *r*_*ℓ*,*π*_(*S*) = (*r*_0_,…, *r*_*ℓ* − 1_).

Recall that the *k*-mers in the vector *h*_*ℓ*,*π*_(*S*) are listed in the order in which they appear in *S*. The order vector *r*_*ℓ*,*π*_ is a permutation of the indices [*ℓ*] that can reorder the *k*-mers according to *π*. That is, *h*_*ℓ*,*π*_(*S*) = (*m*_0_,…, *m*_*ℓ* − 1_) and *i* < *j* imply that 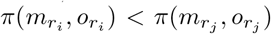 (where, as in the definition, *o*_*i*_ is the occurence number of the *k*-mer *m*_*i*_). The total space usage of a sketch is *O*(log|*S*| +*mℓ*(*k* log *σ*+log *ℓ*)).

The reason for the order vector in the sketch is to recover both an estimate of the weighted Jaccard between the two sequences and how well these common *k*-mers properly align. rore precisely, given two sketches for *S*_1_ and *S*_2_, the number of collisions *h*_*ℓ*,*π*_(*S*_1_) = *h*_*ℓ*,*π*_(*S*_2_) and the number of collisions in the reordered *k*-mers according to the order vector *o*_*ℓ*,*π*_ give an estimate of the weighted Jaccard J^w^(*S*_1_, *S*_2_). Using this estimate, the sizes of *S*_1_ and *S*_2_ from the sketches, and the formula 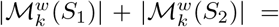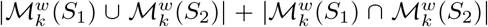, we can recover estimates for the size of the intersection and union of the weighted *k*-mer sets. Finally, formula (10) and (11) give the probability for *ℓ k*-mers from the intersection to be in the same alignment order between the two sequences. The case where the Jaccard similarity is not sufficient to assess that the sequences have a high edit similarity is precisely the case when this last probability is low.

In other words, the extra *O*(log *ℓ*) bits of information per *k*-mer in the OMH sketch compared to a weighted minHash sketch gives corresponds to the supplemental information given by OMH compared to minHash. Given that *ℓ* is small in practice, the cost for this extra information is also very small.

For genomics sequences, it is traditional to compute the minHash using “canonical” *k*-mers (defined as a *k*-mer or its reverse complement, which ever comes first lexicographically). In the OMH sketches, it is not possible to use canonical *k*-mers as this in incompatible with the order information encoded in the vector *h*_*ℓ*,*π*_(*S*). Rather, two sketches, one for the forward strand and one for the reverse are stored. Comparing two sequences requires doing 4 sketches comparisons.

### 4.4 Weighted Jaccard and OMH

Even though the Jaccard and minHash sketches are regularly used as a measure of the *k*-mer content similarity in computational biology software, the weighted Jaccard similarity has been heavily studied and used in other contexts, such as large database document classification and retrieval (e.g., Manasse **et al.**, 2010; Shrivastava, 2016; Wu **et al.**, 2017), near duplicate image detection (Chum **et al.**, 2008), duplicate news story detection (Alonso **et al.**, 2013), source code deduplication (Markovtsev and Kant, 2017), time series indexing (Luo and Shrivastava, 2017), hierarchical topic extraction (Gollapudi and Panigrahy, 2006), or malware classifcation (Drew **et al.**, 2017) and detection (Raff and Nicholas, 2017).

The weighted Jaccard, compared to the unweighted Jaccard, gives a more complete measure of the similarity between two sets or sequences. Obviously, when no elements are repeated, the two similarity are equal. On the other hand, in the case of many repeated elements, the difference can be significant.

For example, returning to the example from Section 2.2 where *S*_1_ = 0…01…1 with *n* − *k* 0s followed by *k* 1s and *S*_2_ with *k* 0s followed by *n − k* 1s, the edit similarity is very low: E_s_(*S*_1_, *S*_2_) ≤ 2*k/n*. The Jaccard similarity is J(*S*_1_, *S*_2_) = 1, in other words, these two sequences are indistinguishable according the Jaccard similarity. On the other hand, the weighted Jaccard is also very low: J^w^(*S*_1_, *S*_2_) = 1/(*n − k*), much more similar to the edit similarity.

In the case of two de Bruijn sequences that might have very low edit similarity, the Jaccard and weighted Jaccard are both equal to 1, as every *k*-mer occurs exactly once. Therefore, in this case the weighted Jaccard provides no extra information. The OMH sketching method, being also sensitive to the relative orders of the *k*-mers (see eq (10)), would have a probability of hash collision much lower than 1.

In Figure 4, we generated one million random binary sequences (*σ* = 2) of length *n* = 100. Each string is then randomly mutated a random number of times (up to 100 times) to obtain a pair of sequences with a random edit distance. Then, for each pair, we compute the actual edit distance, the exact — i.e., not estimated by minHash — Jaccard and weighted Jaccard similarities. Additionally, the OMH sketch (with *ℓ* = 2 and *m* = 500) is also computed for each pair. The graph shows the median and first quartiles computed over the million pairs of sequences. Even for sequences with high edit distance (> 0.4), the Jaccard similarity remains very high. However, the weighted Jaccard and OMH are more sensitive to the edit distance.

**Figure 4.**
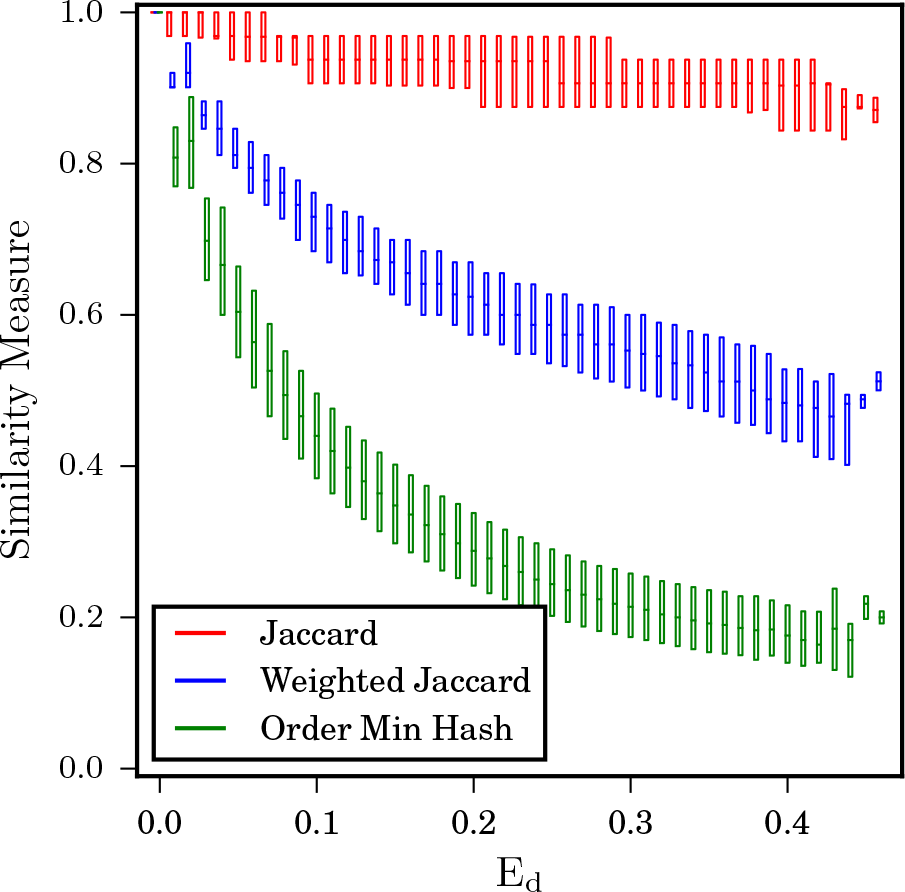
Evolution of the Jaccard, weighted Jaccard and OMH against the edit distance on randomly generated binary sequences. In average, the Jaccard similarity stays high, even for sequences with high edit distance, unlike the weighted Jaccard or OMH which are much more sensitive to the edit distance.

## 5 Conclusion

We presented the OMH method that is an LSH for the edit distance. Unlike the Jaccard similarity, which is only sensitive to the *k*-mer content of a sequence, OMH additionally takes into account the relative order of the *k*-mers in a sequence.

The OMH method is a refinement of the weighted Jaccard similarity that is used extensively in many related fields, such as document classification and duplicate detection. However, despite the advantages of the weighted Jaccard similarity, it has not yet been widely adopted by the bioinformatics community. Using weighted Jaccard and OMH for estimating edit similarity in bioinformatics applications can help reduce the number of false-positive matches which can in-turn avoid unnecessary computations.

## Acknowledgments

The authors would like to thank Natalie Sauerwald for comments on the manuscript.

## References

Aldous, D. and Diaconis, P. (1999). Longest increasing subsequences: From patience sorting to the Baik-Deift-Johansson theorem. Bulletin of the American Mathematical Society, 36(4), 413–432.

Alonso, O., Fetterly, D., and Manasse, M. (2013). Duplicate news story detection revisited. In Asia Information Retrieval Symposium, pages 203–214. Springer.

Backurs, A. and Indyk, P. (2015). Edit distance cannot be computed in strongly subquadratic time (unless SETH is false). In Proceedings of the Forty-Seventh Annual ACM Symposium on Theory of Computing, STOC ’15, pages 51–58, New York, NY, USA. ACM.

Bar-Yossef, Z., Jayram, T. S., Krauthgamer, R., and Kumar, R. (2004). Approximating edit distance efficiently. In 45th Annual IEEE Symposium on Foundations of Computer Science, pages 550–559.

Berlin, K., Koren, S., Chin, C.-S., Drake, J. P., Landolin, J. M., and Phillippy, A. M. (2015). Assembling large genomes with single-molecule sequencing and locality-sensitive hashing. Nature Biotechnology, 33(6), 623–630.

Broder, A. Z. (1997). On the resemblance and containment of documents. In Proceedings. Compression and Complexity of SEQUENCES 1997 (Cat. No.97TB100171), pages 21–29.

Chum, O., Philbin, J., Zisserman, A., et al. (2008). Near duplicate image detection: min-hash and tf-idf weighting. In BMVC, volume 810, pages 812–815.

Drew, J., Hahsler, M., and Moore, T. (2017). Polymorphic malware detection using sequence classification methods and ensembles. EURASIP Journal on Information Security, 2017(1), 2.

Fredman, M. L. (1975). On computing the length of longest increasing subsequences. Discrete Mathematics, 11(1), 29–35.

Gollapudi, S. and Panigrahy, R. (2006). Exploiting asymmetry in hierarchical topic extraction. In Proceedings of the 15th ACM international conference on Information and knowledge management, pages 475–482. ACM.

Hunt, J. W. and Szymanski, T. G. (1977). A fast algorithm for computing longest common subsequences. Communications of the ACM, 20(5), 350–353.

Indyk, P. and Motwani, R. (1998). Approximate Nearest Neighbors: Towards Removing the Curse of Dimensionality. In Proceedings of the Thirtieth Annual ACM Symposium on Theory of Computing, STOC ’98, pages 604–613, New York, NY, USA. ACM.

Jaffe, D. B., Butler, J., Gnerre, S., Mauceli, E., Lindblad-Toh, K., Mesirov, J. P., Zody, M. C., and Lander, E. S. (2003). Whole-genome sequence assembly for mammalian genomes: Arachne 2. Genome Research, 13(1), 91–96.

Jain, C., Dilthey, A., Koren, S., Aluru, S., and Phillippy, A. M. (2017). A fast approximate algorithm for mapping long reads to large reference databases. In S. C. Sahinalp, editor, Research in Computational Molecular Biology, pages 66–81, Cham. Springer International Publishing.

Kushilevitz, E., Ostrovsky, R., and Rabani, Y. (2000). Efficient search for approximate nearest neighbor in high dimensional spaces. SIAM Journal on Computing, 30(2), 457–474.

Langmead, B. and Salzberg, S. L. (2012). Fast gapped-read alignment with Bowtie 2. Nature Methods, 9(4), 357–359.

Levenshtein, V. I. (1966). Binary codes capable of correcting deletions, insertions, and reversals. In Soviet Physics Doklady, volume 10, pages 707–710.

Li, H. and Durbin, R. (2010). Fast and accurate long-read alignment with Burrows–Wheeler transform. Bioinformatics, 26(5), 589–595.

Liu, C.-M., Wong, T., Wu, E., Luo, R., Yiu, S.-M., Li, Y., Wang, B., Yu, C., Chu, X., Zhao, K., Li, R., and Lam, T.-W. (2012). SOAP3: ultra-fast GPU-based parallel alignment tool for short reads. Bioinformatics (Oxford, England), 28(6), 878–879.

Luo, C. and Shrivastava, A. (2017). SSH (sketch, shingle, & hash) for indexing massive-scale time series. In NIPS 2016 Time Series Workshop, pages 38–58.

Manasse, M., McSherry, F., and Talwar, K. (2010). Consistent weighted sampling. Unpublished technical report) http://research.microsoft.com/en-us/people/manasse.

Marçais, G., Delcher, A. L., Phillippy, A. M., Coston, R., Salzberg, S. L., and Zimin, A. (2018). MUMmer4: a fast and versatile genome alignment system. PLOS Computational Biology, 14(1), e1005944.

Markovtsev, V. and Kant, E. (2017). Topic modeling of public repositories at scale using names in source code. https://arxiv.org/abs/1704.00135.

Myers, E. W., Sutton, G. G., Delcher, A. L., Dew, I. M., Fasulo, P., Flanigan, M. J., Kravitz, S. A., Mobarry, C. M., Reinert, K. H. J., Remington, K. A., Anson, E. L., Bolanos, R. A., Chou, H.-H., Jordan, C. M., Halpern, A. L., Lonardi, S., Beasley, M., Brandon, R. C., Chen, L., Dunn, P. J., Lai, Z., Liang, Y., Nusskern, D. R., Zhan, M., Zhang, Q., Zheng, X., Rubin, G. M., Adams, M. D., and Venter, J. C. (2000). A whole-genome assembly of Drosophila. Science, 287(5461), 2196–2204.

Ondov, B. D., Treangen, T. J., Melsted, P., Mallonee, A. B., Bergman, N. H., Koren, S., and Phillippy, A. M. (2016). Mash: Fast genome and metagenome distance estimation using MinHash. Genome Biology, 17, 132.

Ostrovsky, R. and Rabani, Y. (2007). Low distortion embeddings for edit distance. J. ACM, 54(5).

Raff, E. and Nicholas, C. (2017). Malware classification and class imbalance via stochastic hashed LZJD. In Proceedings of the 10th ACM Workshop on Artificial Intelligence and Security, pages 111–120. ACM.

Shrivastava, A. (2016). Simple and efficient weighted minwise hashing. In Advances in Neural Information Processing Systems, pages 1498–1506.

van Aardenne-Ehrenfest, T. and de Bruijn, N. (1951). Circuits and trees in oriented linear graphs. Simon Stevin: Wis-En Natuurkundig Tijdschrift, 28, 203–217.

Wu, W., Li, B., Chen, L., and Zhang, C. (2017). Consistent weighted sampling made more practical. In Proceedings of the 26th International Conference on World Wide Web, WWW’17, pages 1035–1043, Republic and Canton of Geneva, Switzerland. International World Wide Web Conferences Steering Committee.

Zhao, M., Lee, W.-P., Garrison, E. P., and Marth, G. T. (2013). SSW Library: An SIMD Smith-Waterman C/C++ Library for Use in Genomic Applications. PLoS ONE, 8(12), e82138.

